# PaSTA: Fast parametric inference of significance for spatial associations between brain maps

**DOI:** 10.64898/2026.03.08.710410

**Authors:** Yuanzhe Liu, Andrew Zalesky

## Abstract

Testing for spatial correlations between pairs of brain maps has emerged as a central task in neuroimaging studies. Determining the statistical significance of such correlations is challenging due to the presence of spatial autocorrelation in brain maps. Here we establish a novel parametric method, Parametric Spatial Test for Associations (PaSTA), to infer the significance of spatial associations between brain maps via covariance-variance modelling and effective degrees of freedom estimation. Our method is fast, reliable, and sensitive, enabling flexible significance testing of brain map correlations over arbitrary cortical surface and brain volumetric domains. We examine the sensitivity and specificity of PaSTA using simulated datasets with known ground truth and demonstrate its utility when applied to empirical brain maps. We extend PaSTA to approximately model modest nonstationarity in spatial autocorrelation and show that, our method yields improved false positive control and statistical power relative to existing approaches when brain maps are spatially heterogeneous.

## Introduction

The last decade has witnessed a rapid proliferation of diverse high-resolution brain maps, facilitated by large consortia and open data-sharing initiatives (Sudlow et al., 2015; Van Essen et al., 2013). Maps derived from histology (Amunts et al., 2013; Paquola et al., 2019), molecular assays (Arnatkevic̆iūtė et al., 2019; Siletti et al., 2023), and multimodal spatiotemporal neuroimaging (Hansen et al., 2022; Margulies et al., 2016; Markello et al., 2022) are now openly available for large cohorts, opening unprecedented opportunities to characterize the spatial, network and hierarchical organization of the brain across scales (Hansen & Misic, 2025).

Quantifying the extent to which a pair of brain maps spatially correspond is a core task in brain mapping. Through spatial comparison of diverse brain maps, organizational principles common across genetics, structure, and functional specialization can be identified in health and disease (Fulcher et al., 2021; Hettwer et al., 2022; Zhang et al., 2025). Spatial correspondence also serves as a straightforward rubric for model evaluation in computational neuroscience, where model-derived maps are considered more plausible when they better recapitulate empirical spatial patterns (D. Li et al., 2025). These applications have made spatial association tests a widely used tool in neuroimaging research.

Spatial autocorrelation is a critical concern when testing for correlations between a pair of brain maps. Nearby measurements are similar and dependent, which violates the independence assumption of most statistical measures of association, reducing the effective degrees of freedom, and leading to inflated false positives, if appropriate corrections are not conducted (Jeganathan et al., 2025; Markello & Misic, 2021).

To address this challenge, methods such as the spin test, BrainSMASH, and eigenstrapping have been developed to estimate the statistical significance of spatial correlations between pairs of brain maps, while accounting for spatial dependencies (Alexander-Bloch et al., 2018; Burt et al., 2020; Koussis et al., 2025). These permutation-based methods are non-parametric and rely on repeated sampling of a suitably defined null distribution for the correlation coefficient. While non-parametric methods can approximate the exact null distribution under certain circumstances, this can be computationally demanding when each permutation involves an expensive operation. This can become intractable when analyses involve statistical inference between multiple pairs of brain maps. Moreover, the accuracy with which surrogate maps produced by current methods adequately handle the challenges of spatial autocorrelation remains to be tested (Bazinet et al., 2025). Other limitations and outstanding issues are discussed elsewhere (Burt et al., 2020; Koussis et al., 2025; Markello & Misic, 2021).

Spatial nonstationarity is increasingly recognized as an important and often unaccounted consideration when conducting statistical inference on brain maps. It refers to spatial variation in the extent of autocorrelation and other statistical properties (e.g., variance and mean) (Bazinet et al., 2025; Hansen & Misic, 2025; Koussis et al., 2025; Leech et al., 2024; Moodie et al., 2024; Scholz et al., 2024). Mounting evidence suggests that when data are nonstationary, established methods may fail to adequately control false positive rates (Koussis et al., 2025; Leech et al., 2024). Accounting for spatial nonstationarity is computationally challenging, and it is unclear whether current methods can effectively and efficiently infer the statistical significance for spatial correspondence between brain maps in the presence of nonstationarity.

Here, we introduce Parametric Spatial Test for Associations (PaSTA) to assess the statistical significance for measures of spatial similarity between pairs of brain maps. PaSTA aims to estimate covariance structures using variograms and infer effective degrees of freedom, enabling parametric null hypothesis testing for correlation coefficients between autocorrelated brain maps. We demonstrate that PaSTA can be straightforwardly extended to approximately model the impact of nonstationarity on degrees of freedom, yielding improved false positive control and statistical power in the presence of spatial heterogeneities compared to established methods. PaSTA also offers flexibility to assess spatial similarity within circumscribed regions of interest, as well as surface and volumetric representations of the whole brain. We present a range of simulations demonstrating that PaSTA is a reliable, fast, and sensitive method to estimate p-values for spatial correlation coefficients between pairs of brain maps, complementing established non-parametric methods that generate nulls through resampling, smoothing, and frequency-superposition. We also demonstrate the novel insights that PaSTA can reveal when applied to statistical significance testing of spatial correlations between empirical brain maps.

## Results

### Overview of PaSTA

We propose PaSTA to parametrically infer the significance of spatial association tests (i.e., spatial correlation coefficients) conducted on brain maps. A schematic overview of the inferential procedure is shown in Fig.1 (also see Methods). PaSTA comprises two steps: First, the covariance structure and spatial dependency within each autocorrelated map is estimated (Fig. 1A-C). To this end, empirical variograms (Burt et al., 2020) are utilized to characterize the variability between data points (i.e., the semivariance) as a function of their spatial separations (i.e., lag distances). Variogram models (Montero et al., 2015) are then fitted to the empirical estimates to obtain a valid, continuous representation for distance-dependent semivariance (Fig. 1B). Covariance matrices are constructed by evaluating the fitted variogram model at all pairwise distances between data points (Fig. 1C).

**Figure 1.**
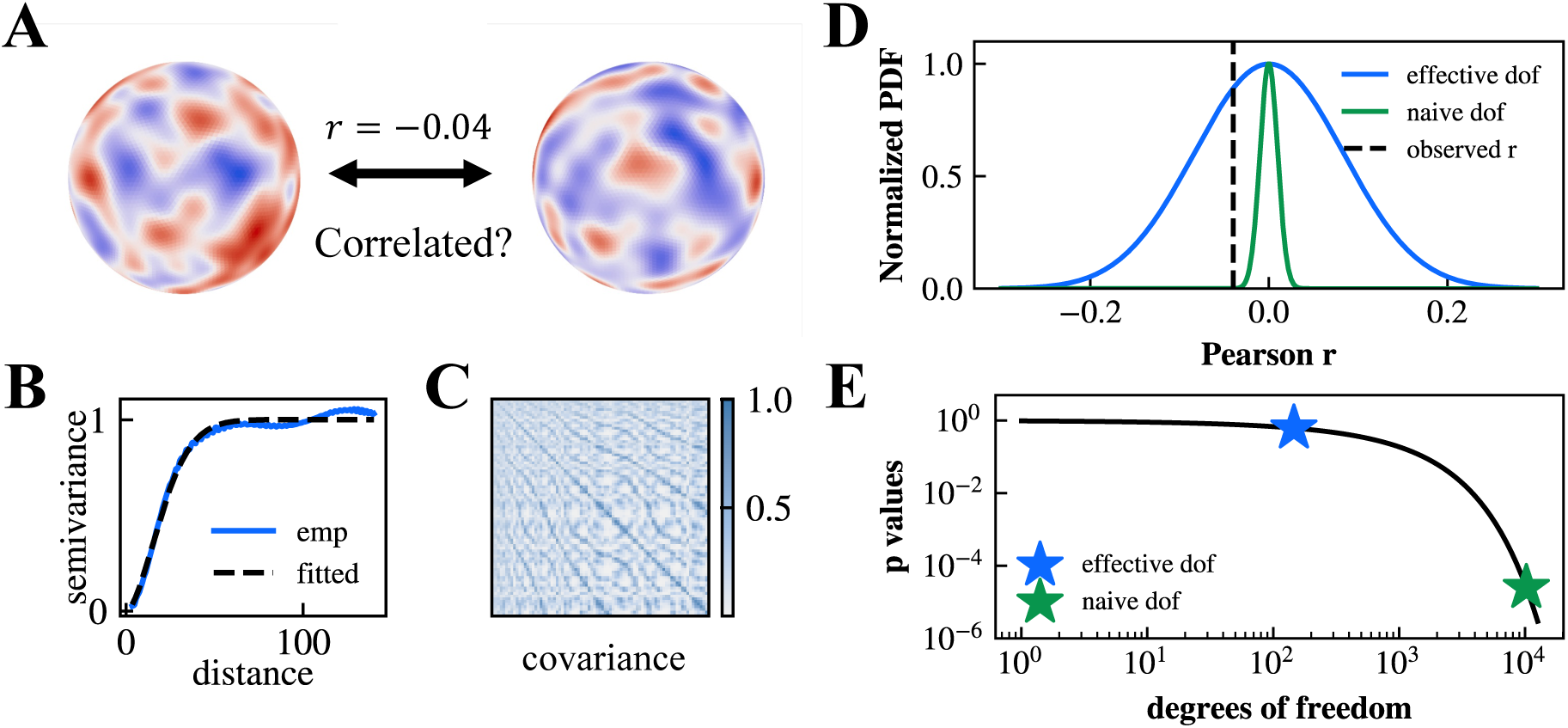
A schematic illustration of PaSTA. A) Example spatial autocorrelated map pair (fsaverage 5 spherical surface, 10,242 vertices) for which PaSTA assesses the significance of Pearson correlation coefficient (*r* = −0.04). B) PaSTA estimates the empirical semivariance (blue) of each map as a function of distance and fits a variogram model (black). C) The covariance matrix of each map is computed from the variogram fit and the distance between observations, which is later used to infer the effective degrees of freedom according to Dutilleul’s derivation. D) Normalized probability density function (PDF) of test statistics (Pearson correlation coefficient) under naïve degrees of freedom (niave dof = 10,240) and effective degrees of freedom (effective dof = 146) computed with PaSTA. The observed Pearson *r* = −0.04 falls in the tail of the distribution under the naive dof, but not under the adjusted effective dof. E) Statistical significance p-value as a function of degrees of freedom for Pearson *r* = −0.04. The null hypothesis is rejected at the significance threshold of *α* = 0.05 when the naive dof is used (*p* = 2.5 × 10^-5^), but it cannot be rejected under the effective dof that accounts for spatial autocorrelation using PaSTA (*p* = 0.62).

Second, the fitted covariance matrices are used to estimate effective degrees of freedom for brain maps, accounting for spatial autocorrelation. Specifically, the effective degrees of freedom are calculated from the covariance matrices using Dutilleul’s derivation (Dutilleul et al., 1993). The significance of the spatial correlation coefficient is finally determined by comparing the observed value against its corresponding parametric null distribution, defined using the estimated effective degrees of freedom (Fig. 1D, 1E).

### Evaluation of statistical power and false positive rate control

We evaluated whether PaSTA controls false positive rates for a nominal statistical significance threshold (*α* = 0.05). To this end, we simulated pairs of independent maps on a spherical mesh (fsaverage5, 10,242 vertices) across a range of spatial autocorrelation strengths (*l* = 0 to 50, ranging from spatial independence to FWHM ≈ 94*mm*, 10,000 map pairs per *l*, Fig. 2A). Due to the independence between all maps, pairs of maps were consistent with the null hypothesis of an absence of spatial correlation, enabling evaluation of false positive rates. We compared the false positive rate control performance of PaSTA to established non-parametric methods, including the spin test and eigenstrapping.

**Figure 2.**
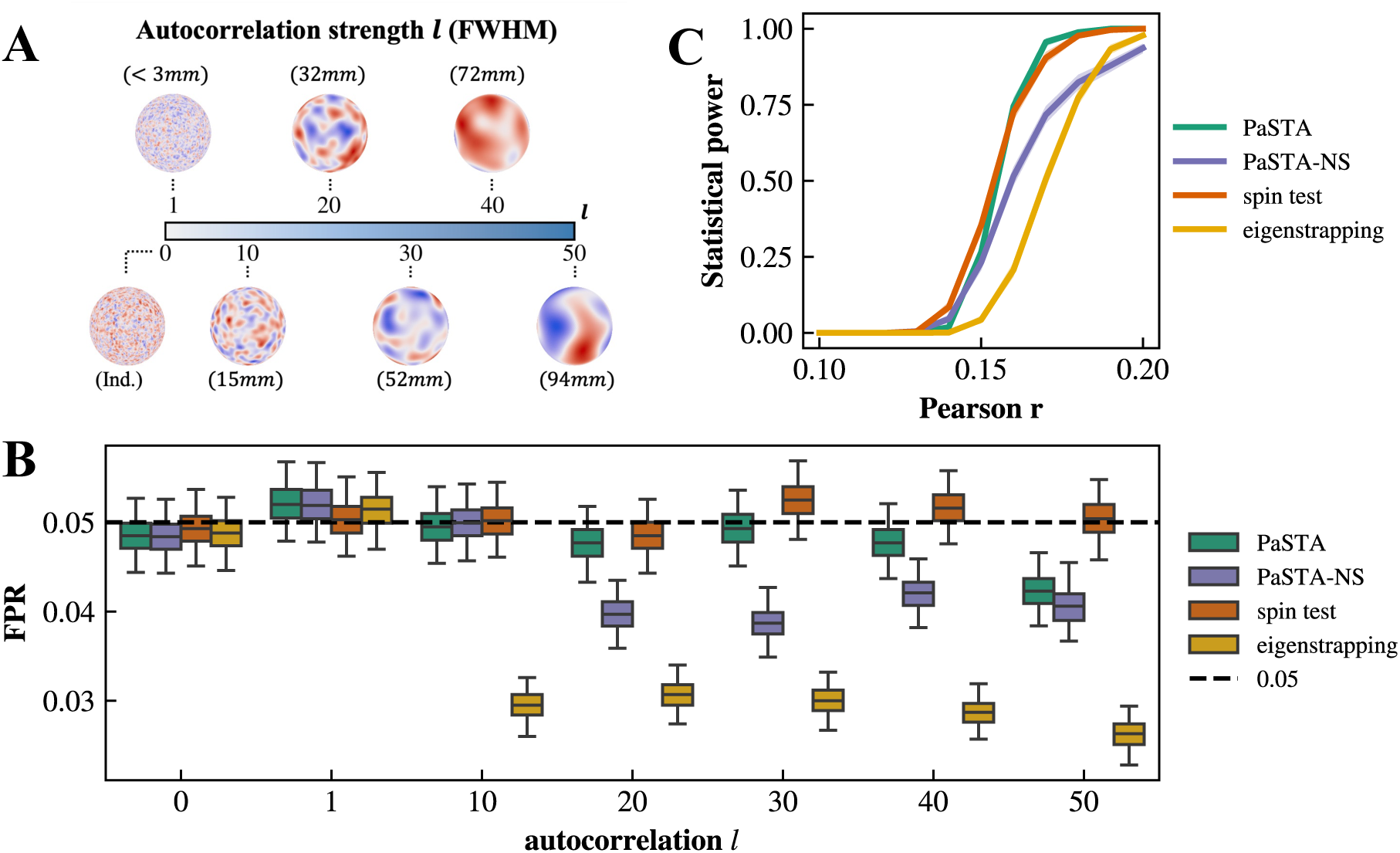
False positive rates and statistical powers of PaSTA and benchmarks. A) False positive rate and statistical power were evaluated with simulated data generated on the spherical brain mesh. Generated data varied in the strength of autocorrelation, ranging from spatial independence to *l* = 50 (FWHM ≈ 94*mm*). B) False positive rates (significance *α* = 0.05) evaluated at all strengths of spatial autocorrelation. Whiskers denote the 95% confidence intervals (CIs) estimated through 1,000 bootstraps. C) Statistical power evaluated at autocorrelation *l* = 20, estimated with 1,000 linearly dependent map pairs at each effect size. Shades indicate 95% CIs.

We found that PaSTA adequately controlled type-I error, observing false positive rates (FPR) at 5% or lower at the significance level of *α* = 0.05 (Fig. 2B). Notably, the spin test maintained an FPR near 5% across all strengths of autocorrelation evaluated, without any evidence of the inflation reported previously (Markello & Misic, 2021). This is expected because autocorrelated maps were generated on the spherical mesh. Data projections between cortical and spherical meshes (Bazinet et al., 2025), which distort autocorrelations in surrogates and lead to increased false positives in the spin test, were not performed in our simulation analysis. PaSTA was mildly overconservative at the strongest autocorrelation evaluated (*l* = 50, FPR of 0.042; 95% CI: [0.0384, 0.0462]). Eigenstrapping was the most conservative approach across all autocorrelation strengths (minimum FPR 0.0263, 95% CI: [0.0232, 0.0296]).

Next, we generated pairs of maps with varying levels of spatial dependency to simulate data consistent with the alternative hypothesis, enabling assessment of statistical power as a function of spatial dependency magnitude, measured with the correlation coefficient. For each magnitude, we generated 1,000 pairs of maps and defined statistical power as the proportion of pairs for which the null hypothesis was rejected (significance *α* = 0.05). Fig. 2C shows statistical power as a function of effect size (also see Fig. S1). These results suggest that PaSTA can provide a viable and fast parametric alternative to current non-parametric methods, achieving comparable performance.

### Modelling spatial nonstationarity with PaSTA-NS

Spatial autocorrelation intrinsic to a brain map may be nonstationary (Leech et al, 2024). This means that a global estimate of autocorrelation may be inaccurate for specific regions of the map where autocorrelation is particularly high or low. If spatial variations in autocorrelation are aligned (i.e., strength of autocorrelation positively correlated) or inversely aligned (i.e., strength of autocorrelation negatively correlated) across the two maps (Fig. 3A), methods that do not account for nonstationarity may either inflate false positive rates or reduce statistical power. When the spatial patterns of autocorrelation in two maps align, the null distribution widens, leading to increased false positives. In contrast, when the spatial patterns inversely align, the null distribution narrows, and true positives are underestimated (Fig. 3B).

**Figure 3.**
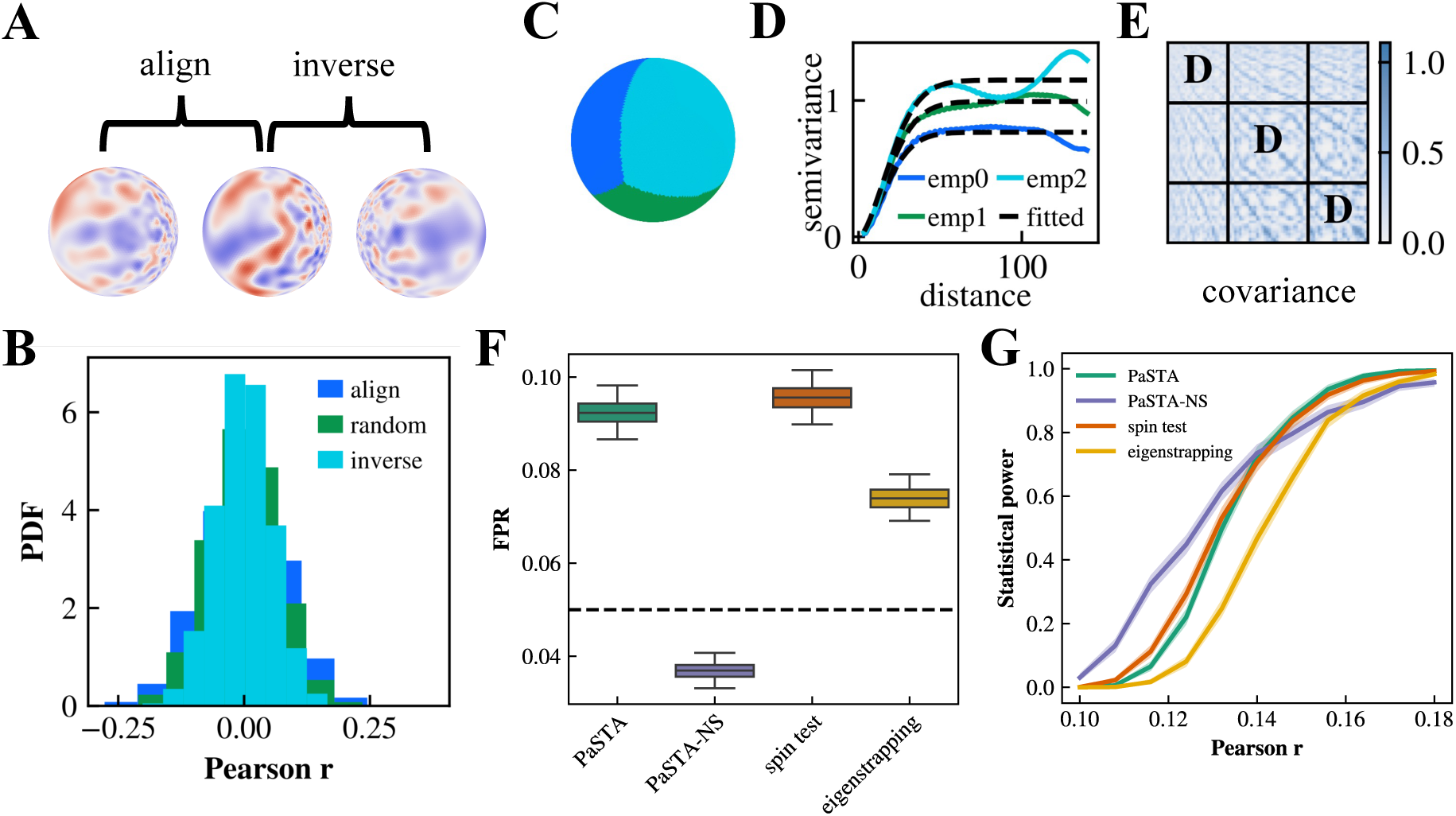
Analyses of FPR and statistical power under nonstationary autocorrelation. A) Example data pairs generated with aligned and inversely aligned autocorrelation. For the aligned pair, both maps show a decreasing autocorrelation from left to right. For the inversely aligned pair, one map shows a decreasing while the other shows an increasing autocorrelation from left to right. B) Alignment and inverse alignment in nonstationary autocorrelation alter the null distribution of Pearson correlation coefficients. C) PaSTA-NS parcellates data to model nonstationarity. An illustrative example with three parcels is displayed. D) PaSTA-NS estimates and fits a separate variogram model for each parcel in C). E) Nonstationary covariance matrix that is constructed by PaSTA-NS. Diagonal blocks (labelled with “D”) stand for within parcel covariance and are computed based on the variogram fit of corresponding parcels. Off-diagonal blocks are inferred using process convolution. F) FPR evaluated with maps that align in nonstationary autocorrelation. G) Statistical power evaluated with maps that inversely align in nonstationary autocorrelation.

To account for the impact of spatial nonstationarity when inferring statistical significance, we developed a generalization of PaSTA called PaSTA-NS (nonstationary). PaSTA-NS aims to approximately model heterogeneity in spatial dependency. This involves estimating covariance-variance structures separately within continuous, non-overlapping parcels derived from spatial clustering (Fig. 3C-E), contrasting to the simpler approach of estimating spatial dependency across the entire spatial domain. Stationary variograms are estimated and fitted within each parcel, which are then integrated into a global nonstationary covariance function using process convolution (Paciorek & Schervish, 2006). As a result, PaSTA-NS approximately accounts for spatial nonstationarity when inferring the effective degrees of freedom. The method is an approximation because nonstationary effects are not necessarily aligned with parcels comprising the spatial clustering.

We evaluated the performance of PaSTA, PaSTA-NS, spin test, and eigenstrapping under a simple model of nonstationary autocorrelation (see Methods). For autocorrelation patterns that are aligned across pairs of brain maps, where FPR control is a concern, independent map pairs were simulated to assess the FPR of each method (Fig. 3F). For inversely aligned autocorrelation, which may reduce statistical power, dependent maps were generated to evaluate the statistical power (Fig. 3G). As expected, inflation in FPR was evident when autocorrelation aligned between the two maps for all methods that did not model nonstationarity. The nonstatationary covariance modeling inherent to PaSTA-NS achieved nominal control of FPR. Importantly, modelling nonstationarity also improved statistical power, reducing the risk of false negative findings arising from inverse alignment in autocorrelation. In contrast, while eigenstrapping reduced false positive findings relative to the spin test and PaSTA, FPR was not controlled and power was reduced compared to the spin test and PaSTA when autocorrelation patterns inversely aligned. We replicated these findings using a different model of nonstationary autocorrelation (Fig. S2) and demonstrated that PaSTA-NS is also effective when (i) data variance of spatial maps is nonstationary (Fig. S2), and (ii) data are spatially homogenous (Fig. 1). These results suggest that PaSTA-NS can approximately mitigate the consequences of nonstationarity, enabling more reliable association inference when data are statistically heterogenous across the space.

### Flexibility across spatial domains

Neuroimaging studies often evaluate dense brain maps that vary in geometry, including surface-based maps and volumetric region-of-interests that may contain missing values (Glasser et al., 2013). As a result, statistical methods that generalize across spatial domains without imposing strict geometric assumptions are useful. For example, it may be of interest to jointly test for spatial correlation across the cortical surface and a volumetric representation of the subcortex. Alternatively, testing for correlation within a circumscribed region of interest may be required. These scenarios are currently challenging with the spin test and eigenstrapping because they would require estimation of a dedicated set of rotations or basis sets that jointly represent both volumetric and surface data, respectively. Together with generative methods such as BrainSMASH (Burt et al., 2020), PaSTA provides a computationally efficient method to perform inference for arbitrary spatial domains and regions of interest.

To demonstrate spatial flexibility, we assessed spatial correlation within: (i) a circumscribed region of interest (ROI) on the sphere (i.e., a quadrant of the spherical mesh comprising 2,528 vertices), and (ii) a cubic volume (20 × 20 × 20, 8,000 voxels). The latter case modelled brain maps represented in the volumetric space. Fig. 4 shows false positive rates under these two spatial domains, assessed using pairs of brain maps consistent with the null hypothesis of absence of spatial correlation within the domain (also see Fig. S3). PaSTA and PaSTA-NS controlled type-I error well across a broad range of autocorrelation strengths in both domains. These results suggest that they provide a reliable, flexible, and efficient approach to test for spatial correspondence in surface- and volume-based neuroimaging studies, enabling region-of-interest and joint cortico-subcortical analyses with and without the presence of missing data.

**Figure 4.**
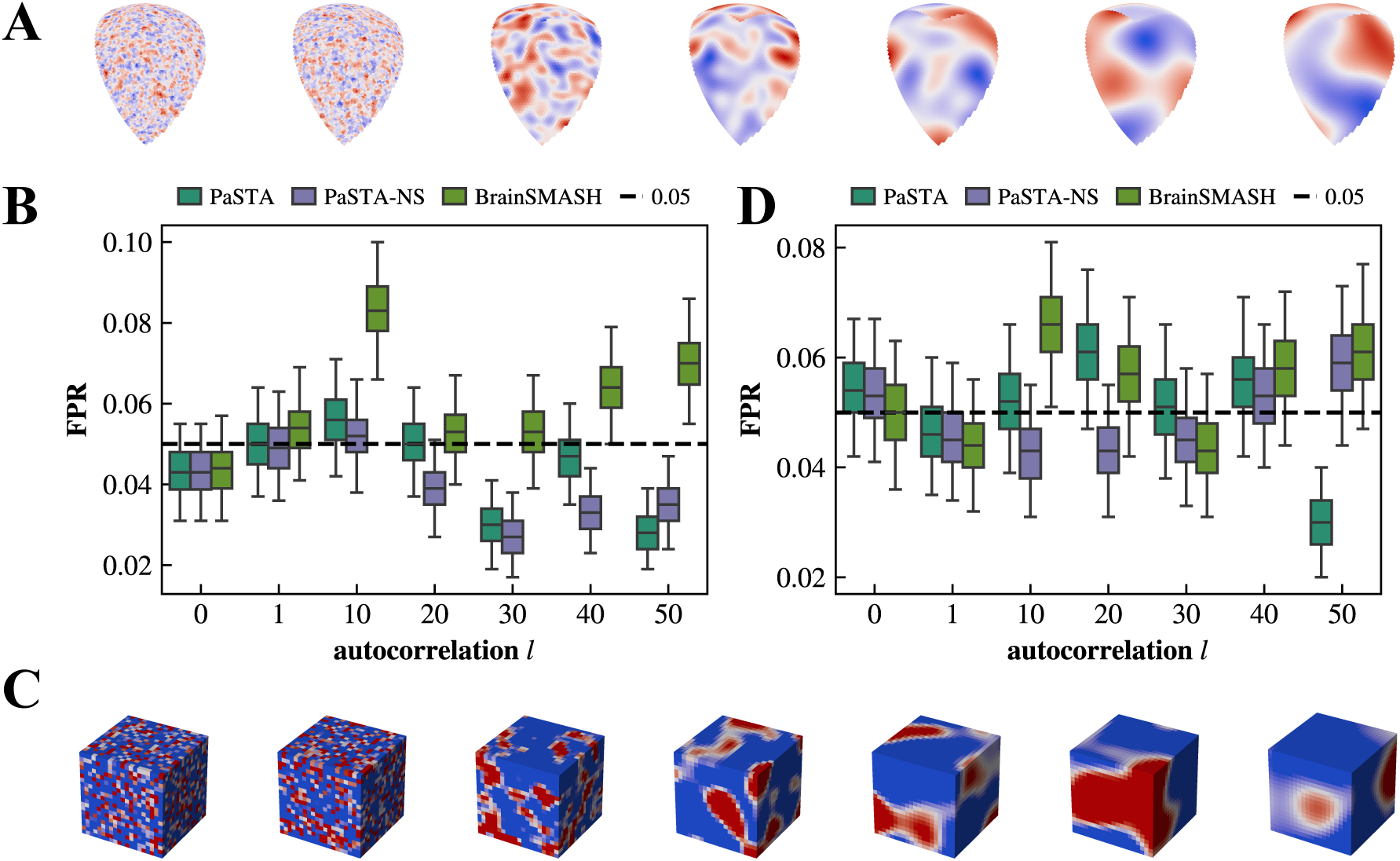
PaSTA controls the nominal false positive rate when evaluating ROI and volumetric data. A) Illustrative example of ROI data simulated on the quadrant of spherical mesh, ranging from spatial independence (left most, *l* = 0) to strong autocorrelation (right most, *l* = 50). B) False positive rate when assessing the simulated data in A), evaluated with 1,000 independent map pairs. C) Illustrative example of simulated volumetric data. Autocorrelation strength increases from *l* = 0 to 50 (left to right). D) Same as B) but assessing the volumetric data in C).

### Assessing correspondence between empirical brain maps

Finally, we demonstrate the use of PaSTA to test for pairwise associations between five brain maps defined on the cortical surface (fsavergae5, 10,242 vertices), including the first principal component of gene expression, Neurosynth, T1/T2 ratio, cortical thickness, and the principal gradient of functional connectivity (Glasser et al., 2013; Margulies et al., 2016; Markello et al., 2021; Yarkoni et al., 2011) (Fig. 5A). These examples span molecular, structural, and functional phenotypes, representing canonical cortical maps (Koussis et al., 2025). Note that low-frequency spatial trends (Bohling, 2005) were first removed from all maps before inference using Laplacian-regularized trend estimation (Barth, 1992). Results for the case without spatial detrending are presented in Fig. S4.

**Figure 5.**
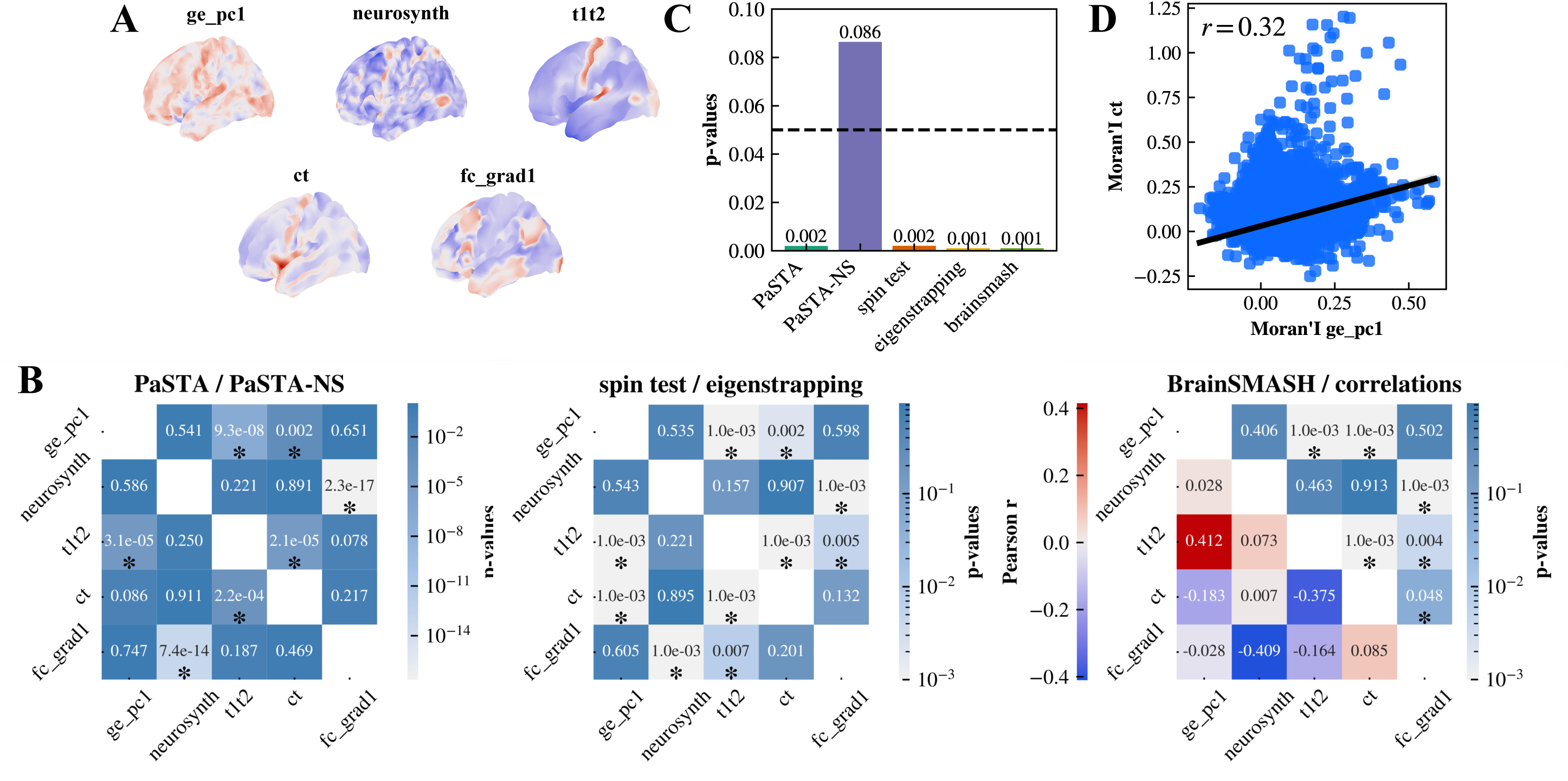
Testing for correlation between empirical brain maps. A) Detrended brain maps evaluated, including the first principal component of gene expression (ge_pc1), neurosynth, T1/T2 ratio (t1t2), cortical thickness (ct), and the principal gradient of functional connectivity (fc_grad1). B) Pairwise associations between empirical brain maps. The matrices display p-values computed using PaSTA (left, upper triangle), PaSTA-NS (left, lower triangle), spin test (center, upper triangle), eigenstrapping (center, lower triangle), and BrainSMASH (right, upper triangle), and the Pearson correlation coefficient (right, lower triangle) for each pair of detrended maps. Significant p-values (<0.05) are marked with the asterisk symbol *. C) Bar plots show p-values for associations between the gene expression and cortical thickness maps. PaSTA-NS found non-significant effects while others yielded significant findings. D) Spatial distributions of autocorrelation are positively correlated between gene expression and cortical thickness maps, measured by local Moran’s I.

PaSTA-NS, PaSTA, spin test, eigenstrapping, and BrainSMASH reported 3, 4, 5, 5, and 6 significant map pairs (significance level 0.05, uncorrected) out of 10 comparisons, respectively (Fig. 5B). PaSTA was more conservative than established non-parametric methods when associating empirical brain maps and rejected the null hypothesis for less pairs of cortical maps. We hypothesize that established methods may have potentially rejected the null hypothesis due to known false positive rate inflation arising from concerns such as spherical projection (Bazinet et al., 2025; Markello & Misic, 2021). However, this requires further testing.

We were particularly interested in the association between gene expression and cortical thickness maps, where PaSTA-NS was unable to reject the null hypothesis (*p* = 0.086), yet other methods including PaSTA reported a significant finding (Fig. 5C). Post-hoc analysis confirmed that the two maps showed positively correlated (i.e., aligned) autocorrelation (measured by local Moran’s I), increasing the likelihood of a false positive finding (Fig. 5D). This suggests that PaSTA-NS may be able to effectively account for data nonstationarity when conducting tests of spatial correlation between real-world brain maps.

## Discussion

Testing for spatial correlation between brain maps is a common task in modern neuroimaging studies. Researchers often seek to determine whether the observed correlation between two maps is statistically significant or instead consistent with the null hypothesis of spatial independence (Arnatkeviciute et al., 2022; Bazinet et al., 2023; Vázquez-Rodríguez et al., 2019; Wan et al., 2025). Numerous non-parametric methods are available to estimate p-values for this null hypothesis. However, non-parametric methods may be slow when testing correlations between many pairs of brain maps and difficult to apply in circumstances where the correlation is computed across a circumscribed region of interest, or when data is represented volumetrically rather than on the cortical surface. These methods may also not offer adequate control of false positive rates in the presence of data nonstationarities (Leech et al., 2024).

To overcome several of these challenges when inferring the statistical significance of correlation coefficients between brain maps, we developed the PaSTA method. PaSTA is the first parametric approximation for this task, complementing existing non-parametric methods such as the spin test, eigenstrapping, and generative approaches (Alexander-Bloch et al., 2018; Burt et al., 2018, 2020; Koussis et al., 2025; Vos De Wael et al., 2020; Weinstein et al., 2021). We provide MATLAB and Python implementations of PaSTA as an open resource for the neuroimaging community (https://github.com/yuanzhel94/PaSTA).

PaSTA involves estimating the covariance structures using established spatial models of semivariance. It enables fast estimation of p-values without needing to generate multiple permutations or surrogates. PaSTA also addresses established concerns with non-parametric methods related to misrepresentation of spatial autocorrelation in surrogates (Bazinet et al., 2025; Markello & Misic, 2021). Surrogate data might not faithfully recapitulate spatial autocorrelation effects due to spherical projection inherent to the spin test, and inappropriate parameter selection in BrainSMASH. While rigorously comparing surrogates with data can mitigate this by refining the data null space, appropriate error control is not guaranteed, and intensive comparison involving many surrogates can introduce additional computational complexity—a concern that does not apply to PaSTA.

The disadvantages of PaSTA are important to acknowledge. The two most important limitations are that: (i) the method assumes data normality; and, (ii) it relies on accurate variogram estimation and fitting. Violation of these assumptions can lead to inflated false positives. However, parametric inference typically remains robust when the violation in data normality is not severe (Knief & Forstmeier, 2021). Furthermore, PaSTA mitigates potential errors in variograms by evaluating semivariance over densely sampled lag distances (Supplementary Fig. S5) and employing a flexible variogram model (i.e., the stable variogram model) that improves model fitting. Importantly, our results suggest that PaSTA provides a comparable or more conservative estimate in empirical brain map analyses relative to established non-parametric benchmarks (Fig. 5).

Spatial nonstationarities in brain maps can alter the null distribution of test statistics and lead to false discoveries. Leech et al. (2024) first discussed false positive rate inflation in the presence of aligned autocorrelation patterns, and this issue has received increasing attention (Bazinet et al., 2025; Hansen & Misic, 2025; Koussis et al., 2025; Scholz et al., 2024). We developed a PaSTA extension called PaSTA-NS to approximately handle the presence of nonstationarities. PaSTA-NS improves false positive control and statistical power when neuroimaging data are spatially heterogenous, without incurring much increase in computational cost compared to PaSTA. This nonstationary generalization provides researchers with an efficient solution for reliable spatial correspondence inference when brain map nonstationarity is evident.

However, PaSTA-NS is an approximation that involves estimating specific variograms within each of many spatial parcels covering the domain over which the correlation is performed. An appropriate parcel size is required for reliable significance inference (see Methods). In addition, nonstationary effects might not align with parcel boundaries, and the choice of parcels can impact the approximation. We determine the parcel size for PaSTA-NS based on the global autocorrelation strength and derive parcels using k-means clustering with spatial coordinates. Parcels can also be defined based on structural and functional priors, such as brain lobes (Fischl, 2012), or an established parcellation atlas. It remains unclear whether nonstationarity is best represented as discrete and circumscribed parcels or a spatial continuum, and this algins with the ongoing discussion between atlas and gradient representations of the brain’s topography (Bernhardt et al., 2022; Glasser et al., 2016; Schaefer et al., 2018). While our simulation demonstrated that parcels can effectively approximate nonstationarity when spatial autocorrelation varies as a continuum (Fig. 3), this approximation is limited up to finite granularity, and future work should investigate the use of PaSTA with parcel-free covariance estimation methods developed more recently (Katzfuss, 2013; Y. Li & Sun, 2019; Zhu & Wu, 2010).

We demonstrated the application of PaSTA to real-world neuroimaging data. We found that PaSTA enables fast estimation of p-values that are broadly consistent with established non-parametric methods. At the same time, discrepancies observed between PaSTA-NS and existing methods in the presence of nonstationary autocorrelation highlight the importance of accounting for spatial heterogeneity when correlating brain maps. Beyond nonstationary autocorrelation, broader forms of spatial heterogeneity including anisotropic autocorrelation, spatial trends, and nonstationary variance, can also cause spurious correlations and lead to false discoveries (Bohling, 2005). These effects have been studied in spatial statistics, yet consideration remains limited in the neuroimaging literature. Further work is needed to investigate these effects.

In summary, PaSTA provides a fast and reliable method to estimate p-values for brain map correlations. We demonstrated the sensitivity and specificity of PaSTA using simulated datasets with known ground truth alongside empirical brain maps. We also showed that PaSTA-NS can enable approximate significance testing in the presence of mild spatial nonstationarities in brain map autocorrelation structure. The main advantages of PaSTA over existing methods are its speed, its ability to handle both volumetric and surface-based maps as well as circumscribed regions of interest, and its capacity to approximately accommodate data nonstationarities.

## Methods

### Estimation of empirical variograms

Let ***x*** = (*X*_1_, …, *X*_*N*_) and ***y*** = (*Y*_1_, …, *Y*_*N*_) denote vectors of observations for the spatial processes *X* and *Y*, respectively, each defined at coordinates (***s***_**1**_, …, ***s***_***N***_). To assess the significance for associations between ***x*** and ***y***, we first utilized variograms to characterize the variability between data points (i.e., semivariance) as a function of their spatial separations (i.e., lag distances). The empirical variogram of ***x*** was estimated broadly following the method described in Burt et al. (2020). Specifically, the empirical semivariance *γ*_*x*_ at lag distance ℎ was computed as

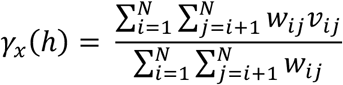

Where 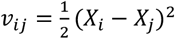 and 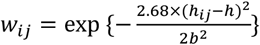 smooths the variogram based on the separation *h*_*ij*_ between observations (*i*, *j*) using a Gaussian kernel with bandwidth *b*. *h*_*ij*_ was defined as 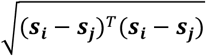 and quantifies Euclidean distances. While not evaluated in the present study, *h*_*ij*_ can be generalized with Mahalanobis distances to account for anisotropic covariance structures. The empirical semivariance was evaluated for *M* lag distances linearly spaced between [*d*_*min*_, *qd* ∗ *d*_*max*_] with interval Δ*h*, where *d*_*min*_ and *d*_*max*_ denote the minimum and maximum distances between observations, and *qd* ∈ (0,1] adjusts the maximum distance evaluated to exclude rare, unreliable estimations for long distance pairs. The bandwidth of Gaussian kernels was set to *b* = 3Δ*h*. Appropriate choices of *M* and *qd* are critical to the accurate estimation of empirical variograms and statistical inference, which is discussed in the Supplementary Materials (Fig. S5). In brief, large *qd* and *M* typically benefits variogram estimation and significance inference, which in theory applies to other variogram-based methods such as BrainSMASH. Throughout this work, we employed *M* = 3 × *sqrt*(*N*) with *N* denoting the number of observations, and *qd* = 0.7. The empirical variogram of ***y*** was estimated following the same procedure.

### Fitting variogram models and constructing covariance matrices

Variogram models were next fitted to empirical variogram estimates. Various variogram models have been proposed in the literature (Chiles & Delfiner, 2012; Matérn, 2013); here, we employed the stable model (Montero et al., 2015):

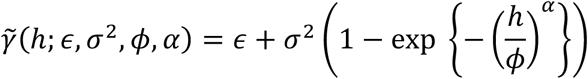

where *ϵ* is the nugget that models the discontinuity at the origin (*h* = 0) and represents microscale variation that the empirical variogram fails to resolve due to finite sampling, *σ*^2^ is the partial sill that measures the semivariance at infinity lag distances (i.e., when data are spatially independent) without considering the nugget, *ϕ* controls the rate that semivariance rises to the sill, and *αϵ*(0,2] determines the smoothness of variograms. Having fitted the variogram models *γ̃*_*x*_ and *γ̃*_*y*_, spatial covariance matrices *C*_*x*_ and *C*_*y*_ were constructed for ***x*** and ***y***, respectively, by evaluating the semivariance between all pairs of observations:

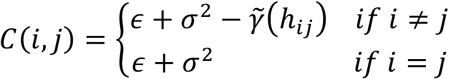

### Inference of effective degrees of freedom and statistical significance

The effective sample size *N*_*ef*_ was computed from covariance matrices *C*_*x*_ and *C*_*y*_ based on Dutilleul’s derivation (Dutilleul et al., 1993)

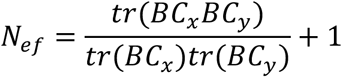

with *B* = *N*^-1^(*I*_*N*_ − *N*^-1^*J*_*N*_), and *I*_*N*_ and *J*_*N*_ denote the *N* × *N* identity matrix and the *N* × *N* matrix of ones, respectively. The adjusted statistical significance is thus computed based on the Pearson correlation coefficient *ρ*(***x***, ***y***) and the effective degrees of freedom (*N*_*ef*_ − 2).

### Parcellations in PaSTA-NS

PaSTA-NS approximates spatial nonstationarity by dividing data into parcels, and therefore its performance depends on the number of parcels derived. Too few, large parcels limit the modeling of fine-scale spatial heterogeneity, while too many, small parcels confine estimation within a limited spatial scope, leading to unreliable variogram estimates and false positives (due to fewer observations, and data points may remain dependent at the largest separation evaluated).

We employed a data-driven approach to define PaSTA-NS parcels that balances heterogeneity detection with reliable variogram estimation. A global variogram was first fitted to all data, and its effective range was computed (the lag distance at which 95% of sill is reached and data are considered effectively independent; defined as 3^1/*α*^*ϕ* for stable variogram models). The number of parcels was then selected such that each parcel approximately spans this effective range from its center, and parcels were subsequently determined via k-means clustering over spatial coordinates of observations. Note that simulations evaluated in the present work span a wide range of autocorrelation strengths. We mandate a maximum partition of 10 to avoid over-partitioning at weak autocorrelation and spatial independence while effectively modelling nonstationarity. We also set a minimum partition of 4 (for spherical meshes, 2 for ROIs and volumes) that prevents PaSTA-NS from collapsing to PaSTA at strong autocorrelations, to assess the impact of parcellations in simulation studies. This latter constraint on minimum partition was not applied when analyzing empirical data.

### Nonstationary covariance function

A variety of nonstationary variogram models have been proposed to estimate and construct spatially varying covariance structures, which can in principle be used in PaSTA-NS to account for data nonstationarity. Here, we employed the process convolution model developed by Paciorek and Schervish (2006). Parcellating spatial data into nonoverlapped groups, stationary local stable variograms (with a common shape parameter *α* that is estimated with the global variogram) were estimated and fitted for each group as described above. Next, these local covariances were integrated into a global nonstationary covariance structure, according to

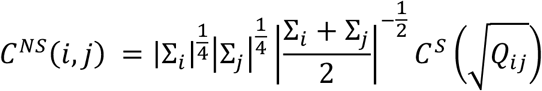

where Σ_*i*_ and Σ_*j*_ are the covariance matrices of the Gaussian kernel centered at point *i* and *j*, respectively, 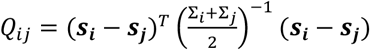, and *C*^*S*^ represents the stationary covariance function. Effective degrees of freedom that account for spatial nonstationarity were subsequently computed according to Dutilleul’s derivation from nonstationary covariance matrices of the two maps, which were further used to infer the statistical significance in PaSTA-NS.

### Data simulation

All simulated data with stationary autocorrelation evaluated in the present work were generated with the GSTools python package (Müller et al., 2022), as described in Bazinet et al. (2025). In brief, autocorrelated maps were generated based on a Gaussian variogram model using the randomization method. Data that varied in autocorrelation strengths were generated with length parameter *l* ranges from 1 to 50, with larger *l* indicates stronger autocorrelation. FWHM of generated maps were estimated using the connectome workbench *wb_command* (Van Essen et al., 2013). In addition, data without spatial autocorrelation (indicated by *l* = 0 throughout the work) were sampled from a standard Gaussian distribution. We simulated data on the fsaverage5 spherical mesh of the brain left hemisphere, a quarter-spherical mesh (fsaverage5 vertices with positive coordinates in the first two axes, 2,528 vertices), and a cubic lattice with 20^B^ voxels (edge length = 5*mm*). False positive rates were evaluated with independently generated map pairs, computed as the probability of independent map pairs being significantly associated at a significance level of *α* = 0.05. Statistical power was estimated with linearly related data that vary in effective sizes (measured by Pearson correlation coefficient), computed as the probability of related map pairs being significantly associated at a significance level of *α* = 0.05. Unless otherwise specified, FPR and statistical power were evaluated with 10,000 and 1,000 map pairs, respectively.

To simulate data with nonstationary autocorrelation, we first drew spatially independent samples from the standard Gaussian distribution. Nonstationary autocorrelation was introduced by applying Gaussian kernel smoothing, with kernel bandwidths linearly varying along the first axis of spatial coordinates. We reported kernel bandwidths varying between [1, 20]mm in the main, and between [1, 30]mm in Supplementary Materials. Specifically, independent map pairs with aligned autocorrelation patterns were used to assess false positive rates, whereas correlated pairs with inversely aligned autocorrelation patterns were generated to evaluate statistical power. Linear dependence between correlated pairs was introduced prior to kernel smoothing.

### Non-parametric methods

Established non-parametric methods tested included the spin test, eigenstrapping, and BrainSMASH (Alexander-Bloch et al., 2018; Burt et al., 2020; Koussis et al., 2025). All these methods evaluate the statistical significance by generating a null distribution of autocorrelation-preserved surrogate maps, and we generated 1,000 surrogates in each run. Spin test and eigenstrapping were selected for simulations on the spherical brain mesh because both methods are inherently linked to the spherical geometry. Specifically, the spin test generates surrogates by randomly rotating data on the spherical mesh. Eigenstrapping constructs surrogates through random rotation of geometric eigenmodes, which are equivalent to spherical harmonics on the spherical mesh. Choices of amplitude adjustment, residual treatment, and number of modes retained in eigenstrapping can affect surrogate quality and Type-I error control; we performed amplitude adjustment without adding residuals back and optimized the number of modes for each map as suggested in the original work. All eigenmodes with spatial wavelength longer than 2 × *FWHM* of data were retained, where FWHMs were estimated using the connectome workbench *wb_command* (Van Essen et al., 2013). BrainSMASH were chosen to benchmark analyses on the mesh ROI and cubic volume because of its flexibility across spatial geometries. Parameter choices in BrainSMASH are critical to its performance—larger number of distance bins and broader range of distances (i.e., *nh* and *pv* parameters in its python implementation) are expected to improve false positive control, as discussed in the variogram estimation section above and Supplementary Materials. However, running large-scale BrainSMASH simulations on dense spatial maps with large *nh* and *pv* parameters is computationally unaffordable. Therefore, we adopted a balanced setting with *nh* = 50 and *pv* = 25 when autocorrelation was weak (*l* ≤ 10), and *nh* = 25 and *pv* = 70 when autocorrelation was strong (*l* ≥ 20). For BrainSMASH, FPRs were evaluated using 1,000 simulated map pairs.

### Brain map detrending

Empirical brain maps show spatial trends—large-scale, deterministic, spatial shifts in the mean that fundamentally differs from the local, stochastic spatial autocorrelation (Bohling, 2005). These spatial trends (or nonstationary means) are known to induce spurious correlations.

To account for trend effects when associating empirical brain maps, we decomposed each brain map *X* into the spatial trend *m* and autocorrelated residuals *ε*. This was achieved using Laplacian regularized trend estimation *m* = (*I* + *λK*)^-1^*X*, where *K* is the cotangent Laplacian, and the non-negative *λ* is a regularization parameter to be optimized (Barth, 1992). The detrended residuals *ɛ* is computed by *ɛ* = *X* − *m*. We fitted trends and residuals using *λ* ∈ [10^-1^, 1, 10,10*, 10^B^] and optimized *λ* by inspecting the variograms of residuals and selecting the value corresponding to a stable sill indicative of absence of trends. Associations between brain maps were subsequently evaluated on detrended residuals to account for spatial confounds. We also tested for associations between raw brain maps with spatial trends and reported results in Fig. S4.

## Notes

### Competing Interest Statement

The authors have declared no competing interest.

### Summary of Updates

Name of method changed from SPICE to PaSTA to distinguish from an existing method SPICE developed prior to our work.

## References

Alexander-Bloch, A. F., Shou, H., Liu, S., Satterthwaite, T. D., Glahn, D. C., Shinohara, R. T., Vandekar, S. N., & Raznahan, A. (2018). On testing for spatial correspondence between maps of human brain structure and function. NeuroImage, 178, 540–551. 10.1016/j.neuroimage.2018.05.070

Amunts, K., Lepage, C., Borgeat, L., Mohlberg, H., Dickscheid, T., Rousseau, M.-É., Bludau, S., Bazin, P.-L., Lewis, L. B., Oros-Peusquens, A.-M., Shah, N. J., Lippert, T., Zilles, K., & Evans, A. C. (2013). BigBrain: An Ultrahigh-Resolution 3D Human Brain Model. Science, 340(6139), 1472–1475. 10.1126/science.1235381

Arnatkeviciute, A., Fulcher, B. D., Bellgrove, M. A., & Fornito, A. (2022). Imaging Transcriptomics of Brain Disorders. Biological Psychiatry Global Open Science, 2(4), 319–331. 10.1016/j.bpsgos.2021.10.002

Arnatkevic̆iūtė, A., Fulcher, B. D., & Fornito, A. (2019). A practical guide to linking brain-wide gene expression and neuroimaging data. NeuroImage, 189, 353–367. 10.1016/j.neuroimage.2019.01.011

Barth, T. J. (1992). ASPECTS OF UNSTRUCTURED GRIDS AND FINITE-VOLUME SOLVERS FOR THE EULER AND NAVIER-STOKES EQUATIONS. AGARD.

Bazinet, V., Hansen, J. Y., & Misic, B. (2023). Towards a biologically annotated brain connectome. Nature Reviews Neuroscience, 24(12), 747–760. 10.1038/s41583-023-00752-3

Bazinet, V., Liu, Z.-Q., & Misic, B. (2025). The effect of spherical projection on spin tests for brain maps. Imaging Neuroscience, 3, IMAG.a.118. 10.1162/IMAG.a.118

Bernhardt, B. C., Smallwood, J., Keilholz, S., & Margulies, D. S. (2022). Gradients in brain organization. NeuroImage, 251, 118987. 10.1016/j.neuroimage.2022.118987

Bohling, G. (2005). INTRODUCTION TO GEOSTATISTICS And VARIOGRAM ANALYSIS. Kansas Geological Survey.

Burt, J. B., Demirtaş, M., Eckner, W. J., Navejar, N. M., Ji, J. L., Martin, W. J., Bernacchia, A., Anticevic, A., & Murray, J. D. (2018). Hierarchy of transcriptomic specialization across human cortex captured by structural neuroimaging topography. Nature Neuroscience, 21(9), 1251–1259. 10.1038/s41593-018-0195-0

Burt, J. B., Helmer, M., Shinn, M., Anticevic, A., & Murray, J. D. (2020). Generative modeling of brain maps with spatial autocorrelation. NeuroImage, 220, 117038. 10.1016/j.neuroimage.2020.117038

Chiles, J.-P., & Delfiner, P. (2012). Geostatistics: Modeling spatial uncertainty. John Wiley & Sons.

Dutilleul, P., Clifford, P., Richardson, S., & Hemon, D. (1993). Modifying the t Test for Assessing the Correlation Between Two Spatial Processes. Biometrics, 49(1), 305. 10.2307/2532625

Fischl, B. (2012). FreeSurfer. NeuroImage, 62(2), 774–781. 10.1016/j.neuroimage.2012.01.021

Fulcher, B. D., Arnatkeviciute, A., & Fornito, A. (2021). Overcoming false-positive gene-category enrichment in the analysis of spatially resolved transcriptomic brain atlas data. Nature Communications, 12(1), 2669. 10.1038/s41467-021-22862-1

Glasser, M. F., Coalson, T. S., Robinson, E. C., Hacker, C. D., Harwell, J., Yacoub, E., Ugurbil, K., Andersson, J., Beckmann, C. F., Jenkinson, M., Smith, S. M., & Van Essen, D. C. (2016). A multi-modal parcellation of human cerebral cortex. Nature, 536(7615), 171–178. 10.1038/nature18933

Glasser, M. F., Sotiropoulos, S. N., Wilson, J. A., Coalson, T. S., Fischl, B., Andersson, J. L., Xu, J., Jbabdi, S., Webster, M., Polimeni, J. R., Van Essen, D. C., & Jenkinson, M. (2013). The minimal preprocessing pipelines for the Human Connectome Project. NeuroImage, 80, 105–124. 10.1016/j.neuroimage.2013.04.127

Hansen, J. Y., & Misic, B. (2025). Integrating and interpreting brain maps. Trends in Neurosciences, S0166223625001249. 10.1016/j.tins.2025.06.003

Hansen, J. Y., Shafiei, G., Markello, R. D., Smart, K., Cox, S. M. L., Nørgaard, M., Beliveau, V., Wu, Y., Gallezot, J.-D., Aumont, É., Servaes, S., Scala, S. G., DuBois, J. M., Wainstein, G., Bezgin, G., Funck, T., Schmitz, T. W., Spreng, R. N., Galovic, M., … Misic, B. (2022). Mapping neurotransmitter systems to the structural and functional organization of the human neocortex. Nature Neuroscience, 25(11), 1569–1581. 10.1038/s41593-022-01186-3

Hettwer, M. D., Larivière, S., Park, B. Y., Van Den Heuvel, O. A., Schmaal, L., Andreassen, O. A., Ching, C. R. K., Hoogman, M., Buitelaar, J., Van Rooij, D., Veltman, D. J., Stein, D. J., Franke, B., Van Erp, T. G. M., ENIGMA ADHD Working Group, ENIGMA Autism Working Group, Van Rooij, D., ENIGMA Bipolar Disorder Working Group, ENIGMA Major Depression Working Group, … Valk, S. L. (2022). Coordinated cortical thickness alterations across six neurodevelopmental and psychiatric disorders. Nature Communications, 13(1), 6851. 10.1038/s41467-022-34367-6

Jeganathan, J., Koussis, N. C., Paton, B., Phogat, R., Pang, J., Mansour L, S., Zalesky, A., & Breakspear, M. (2025). Spurious correlations in surface-based functional brain imaging. Imaging Neuroscience, 3, imag_a_00478. 10.1162/imag_a_00478

Katzfuss, M. (2013). Bayesian Nonstationary Spatial Modeling for Very Large Datasets. Environmetrics, 24(3), 189–200. 10.1002/env.2200

Knief, U., & Forstmeier, W. (2021). Violating the normality assumption may be the lesser of two evils. Behavior Research Methods, 53(6), 2576–2590. 10.3758/s13428-021-01587-5

Koussis, N. C., Pang, J. C., Phogat, R., Jeganathan, J., Paton, B., Fornito, A., Robinson, P. A., Misic, B., & Breakspear, M. (2025). Generation of surrogate brain maps preserving spatial autocorrelation through random rotation of geometric eigenmodes. Imaging Neuroscience, 3, IMAG.a.71. 10.1162/IMAG.a.71

Leech, R., Smallwood, J., Moran, R., Jones, E., Vowles, N., Leech, D., Viegas, E., Turkheimer, F., Alberti, F., Margulies, D., Jefferies, E., Alexander-Bloch, A., Zhou, Y., Bernhardt, B., & Váša, F. (2024). The impact of heterogeneous spatial autocorrelation on comparisons of brain maps. Neuroscience. 10.1101/2024.06.14.598987

Li, D., Zalesky, A., Wang, Y., Wang, H., Ma, L., Cheng, L., Banaschewski, T., Barker, G. J., Bokde, A. L. W., Brühl, R., Desrivières, S., Flor, H., Garavan, H., Gowland, P., Grigis, A., Heinz, A., Lemaître, H., Martinot, J.-L., Martinot, M.-L. P., … IMAGEN Consortium. (2025). Mapping the coupling between tract reachability and cortical geometry of the human brain. Nature Communications, 16(1), 7489. 10.1038/s41467-025-62812-9

Li, Y., & Sun, Y. (2019). Efficient Estimation of Non-stationary Spatial Covariance Functions with Application to High-resolution Climate Model Emulation. Statistica Sinica. 10.5705/ss.202017.0536

Margulies, D. S., Ghosh, S. S., Goulas, A., Falkiewicz, M., Huntenburg, J. M., Langs, G., Bezgin, G., Eickhoff, S. B., Castellanos, F. X., Petrides, M., Jefferies, E., & Smallwood, J. (2016). Situating the default-mode network along a principal gradient of macroscale cortical organization. Proceedings of the National Academy of Sciences, 113(44), 12574–12579. 10.1073/pnas.1608282113

Markello, R. D., Arnatkeviciute, A., Poline, J.-B., Fulcher, B. D., Fornito, A., & Misic, B. (2021). Standardizing workflows in imaging transcriptomics with the abagen toolbox. eLife, 10, e72129. 10.7554/eLife.72129

Markello, R. D., Hansen, J. Y., Liu, Z.-Q., Bazinet, V., Shafiei, G., Suárez, L. E., Blostein, N., Seidlitz, J., Baillet, S., Satterthwaite, T. D., Chakravarty, M. M., Raznahan, A., & Misic, B. (2022). neuromaps: Structural and functional interpretation of brain maps. Nature Methods, 19(11), 1472–1479. 10.1038/s41592-022-01625-w

Markello, R. D., & Misic, B. (2021). Comparing spatial null models for brain maps. NeuroImage, 236, 118052. 10.1016/j.neuroimage.2021.118052

Matérn, B. (2013). Spatial variation (Vol. 36). Springer Science & Business Media.

Montero, J.-M., Fernández-Avilés, G., & Mateu, J. (2015). Spatial and spatio-temporal geostatistical modeling and kriging. John Wiley & Sons.

Moodie, J. E., Buchanan, C., Furtjes, A., Conole, E., Stolicyn, A., Corley, J., Ferguson, K., Hernandez, M. V., Maniega, S. M., Russ, T. C., Luciano, M., Whalley, H., Bastin, M. E., Wardlaw, J., Deary, I., & Cox, S. (2024). Brain maps of general cognitive function and spatial correlations with neurobiological cortical profiles. Neuroscience. 10.1101/2024.12.17.628670

Müller, S., Schüler, L., Zech, A., & Heße, F. (2022). GSTools v1.3: A toolbox for geostatistical modelling in Python. Geoscientific Model Development, 15(7), 3161–3182. 10.5194/gmd-15-3161-2022

Paciorek, C. J., & Schervish, M. J. (2006). Spatial modelling using a new class of nonstationary covariance functions. Environmetrics, 17(5), 483–506. 10.1002/env.785

Paquola, C., Vos De Wael, R., Wagstyl, K., Bethlehem, R. A. I., Hong, S.-J., Seidlitz, J., Bullmore, E. T., Evans, A. C., Misic, B., Margulies, D. S., Smallwood, J., & Bernhardt, B. C. (2019). Microstructural and functional gradients are increasingly dissociated in transmodal cortices. PLOS Biology, 17(5), e3000284. 10.1371/journal.pbio.3000284

Schaefer, A., Kong, R., Gordon, E. M., Laumann, T. O., Zuo, X.-N., Holmes, A. J., Eickhoff, S. B., & Yeo, B. T. T. (2018). Local-Global Parcellation of the Human Cerebral Cortex from Intrinsic Functional Connectivity MRI. Cerebral Cortex, 28(9), 3095–3114. 10.1093/cercor/bhx179

Scholz, R., Benn, R. A., Shevchenko, V., Klatzmann, U., Wei, W., Alberti, F., Chiou, R., Zhang, X.-H., Leech, R., Smallwood, J., & Margulies, D. S. (2024). Individual brain activity patterns during task are predicted by distinct resting-state networks that may reflect local neurobiological features. Neuroscience. 10.1101/2024.11.13.621472

Siletti, K., Hodge, R., Mossi Albiach, A., Lee, K. W., Ding, S.-L., Hu, L., Lönnerberg, P., Bakken, T., Casper, T., Clark, M., Dee, N., Gloe, J., Hirschstein, D., Shapovalova, N. V., Keene, C. D., Nyhus, J., Tung, H., Yanny, A. M., Arenas, E., … Linnarsson, S. (2023). Transcriptomic diversity of cell types across the adult human brain. Science, 382(6667), eadd7046. 10.1126/science.add7046

Sudlow, C., Gallacher, J., Allen, N., Beral, V., Burton, P., Danesh, J., Downey, P., Elliott, P., Green, J., Landray, M., Liu, B., Matthews, P., Ong, G., Pell, J., Silman, A., Young, A., Sprosen, T., Peakman, T., & Collins, R. (2015). UK Biobank: An Open Access Resource for Identifying the Causes of a Wide Range of Complex Diseases of Middle and Old Age. PLOS Medicine, 12(3), e1001779. 10.1371/journal.pmed.1001779

Van Essen, D. C., Smith, S. M., Barch, D. M., Behrens, T. E. J., Yacoub, E., & Ugurbil, K. (2013). The WU-Minn Human Connectome Project: An overview. NeuroImage, 80, 62–79. 10.1016/j.neuroimage.2013.05.041

Vázquez-Rodríguez, B., Suárez, L. E., Markello, R. D., Shafiei, G., Paquola, C., Hagmann, P., Van Den Heuvel, M. P., Bernhardt, B. C., Spreng, R. N., & Misic, B. (2019). Gradients of structure–function tethering across neocortex. Proceedings of the National Academy of Sciences, 116(42), 21219–21227. 10.1073/pnas.1903403116

Vos De Wael, R., Benkarim, O., Paquola, C., Lariviere, S., Royer, J., Tavakol, S., Xu, T., Hong, S.-J., Langs, G., Valk, S., Misic, B., Milham, M., Margulies, D., Smallwood, J., & Bernhardt, B. C. (2020). BrainSpace: A toolbox for the analysis of macroscale gradients in neuroimaging and connectomics datasets. Communications Biology, 3(1), 103. 10.1038/s42003-020-0794-7

Wan, B., He, Y., Warrier, V., John, A., Kirschner, M., Eickhoff, S. B., Bethlehem, R. A. I., & Valk, S. L. (2025). Genetic, transcriptomic, metabolic, and neuropsychiatric underpinnings of cortical functional gradients. Psychiatry and Clinical Psychology. 10.1101/2025.03.03.25323242

Weinstein, S. M., Vandekar, S. N., Adebimpe, A., Tapera, T. M., Robert-Fitzgerald, T., Gur, R. C., Gur, R. E., Raznahan, A., Satterthwaite, T. D., Alexander-Bloch, A. F., & Shinohara, R. T. (2021). A simple permutation-based test of intermodal correspondence. Human Brain Mapping, 42(16), 5175–5187. 10.1002/hbm.25577

Yarkoni, T., Poldrack, R. A., Nichols, T. E., Van Essen, D. C., & Wager, T. D. (2011). Large-scale automated synthesis of human functional neuroimaging data. Nature Methods, 8(8), 665–670. 10.1038/nmeth.1635

Zhang, X.-H., Anderson, K. M., Dong, H.-M., Chopra, S., Dhamala, E., Emani, P. S., Gerstein, M. B., Margulies, D. S., & Holmes, A. J. (2025). The cell-type underpinnings of the human functional cortical connectome. Nature Neuroscience, 28(1), 150–160. 10.1038/s41593-024-01812-2

Zhu, Z., & Wu, Y. (2010). Estimation and Prediction of a Class of Convolution-Based Spatial Nonstationary Models for Large Spatial Data. Journal of Computational and Graphical Statistics, 19(1), 74–95. 10.1198/jcgs.2009.07123

