## supplementary_information for "PaSTA: Fast parametric inference of significance for spatial associations between brain maps"

#### Statistical power under stationary autocorrelation that vary in strengths

The FPR of methods for spatial association tests varies depending on autocorrelation strengths, and so is their statistical power. We evaluated the power of PaSTA, PaSTA-NS, spin test, and eigesntrapping using stationary autocorrelated data simulated on the spherical brain mesh, at various strengths of autocorrelations. Results at $l = 20$ were reported in Fig. 2C in the main. Here, we disclosed results of sensitivity at $l \in [0, 1, 10, 30, 40, 50]$ in Fig. S1. Conservative methods (i.e., smaller FPR in Fig. 2B) are typically less sensitive (smaller power).

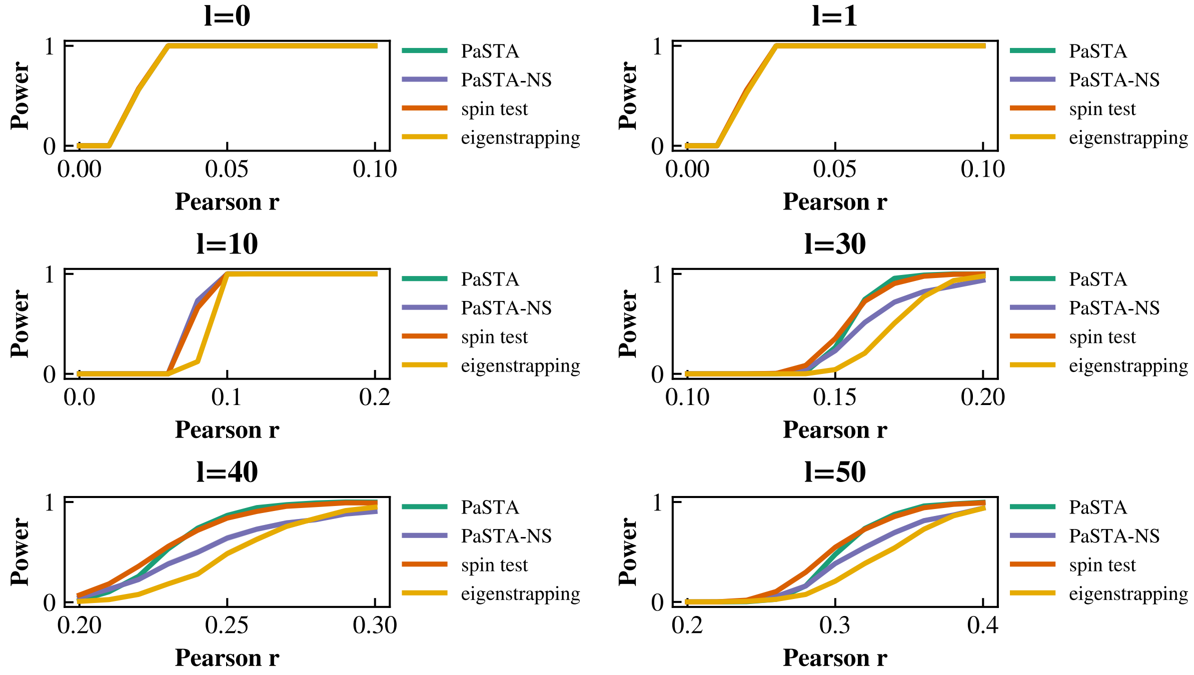


*Figure S1. Statistical power of PaSTA, PaSTA-NS, spin test, and eigenstrapping when assessing simulated maps with stationary autocorrelation.* Data were simulated with autocorrelation strengths ranging from spatial independence ($l = 0$) to strong autocorrelation ($l = 50$).

### PaSTA-NS improves specificity and sensitivity under data nonstationarity

We developed PaSTA-NS to account for nonstationary autocorrelation and variance when assessing map-to-map similarity. Importantly, this improves the specificity when patterns of nonstationarity aligns, and sensitivity under inverse alignment. We demonstrate this with data vary in autocorrelation (Gaussian smoothing kernel bandwidths range from 1 mm to 20 mm) in Fig. 3. Here, we replicate the results using a different extent of spatial nonstationarity in autocorrelation (smoothing kernal bandwidths ranging from 1 mm to 30 mm, Fig. S2A-D), and data simulated with nonstationary variance (standard deviation ranging from 1 to 3, Fig. S2E-H, and from 1 to 4, Fig. S2I-L).

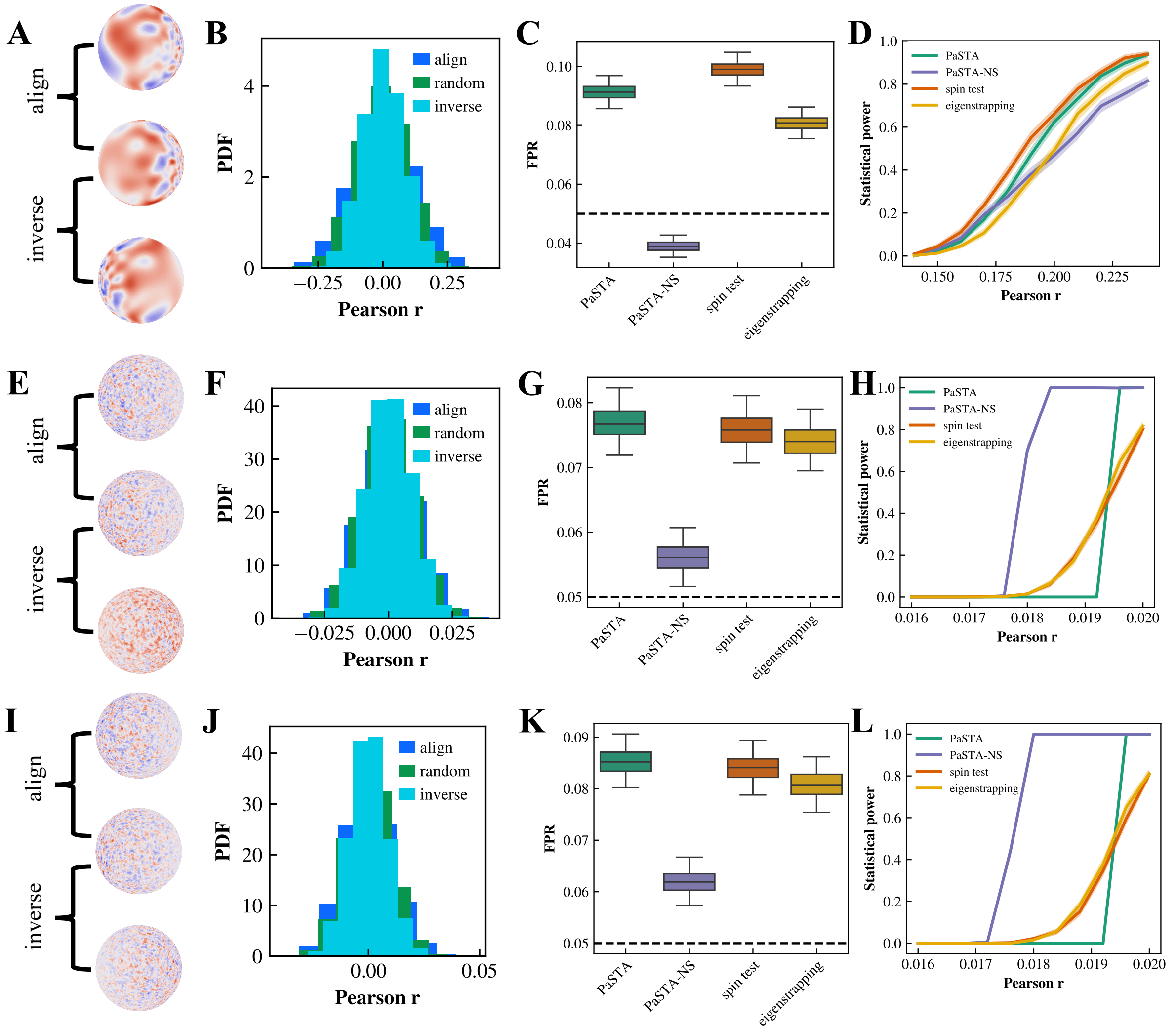


*Figure S2. Replication of modelling nonstationarity with PaSTA-NS. A)-D) Same as Fig. 3 but with a different range of nonstationary autocorrelation (smoothed with kernal bandwidths from 1mm to 30 mm). A) Visualization of example data. B) Alignment and inverse alignment alter null distribution. C) FPR evaluated with aligned pairs. D) Statistical power evaluated with inversely aligned pairs. E)-H) Same as A)-D) but for nonstationary variance changing from standard deviation of 1 to 3. I)-L) Same as A)-D) but for nonstationary variance changing from standard deviation of 1 to 4.*

### FPR of PaSTA over mesh ROI and volumetric data

We demonstrate the geometrical flexibility of PaSTA using data simulated on the mesh ROI and cubic volume in Fig. 4, compared to BrainSMASH. However, due to the computational intensiveness of BrainSMASH, comparison was only conducted for 1,000 independent map pairs per autocorrelation strengths. We further increased the number of simulations to 10,000 map pairs per autocorrelation strengths to provide a more detailed evaluation of PaSTA. As shown in Fig. S3, PaSTA turns mildly overconservative at strong autocorrelation strengths, similar to observations in Fig. 2. Note that for volumetric simulations, PaSTA-NS becomes less conservative than PaSTA at $l = 40$ and 50, and its FPR is higher than 0.05 at $l = 50$ (Fig. S3, right). This is because we forced PaSTA-NS to evaluate data nonstationarity using at least 2 parcels, with the aim to distinguish it from PaSTA in simulation analysis (also see Methods). However, simulated volumetric data at strong autocorrelation strengths show an autocorrelation range longer than the size of half-split parcels, leading to the increase in FPR. When the minimum parcel restriction is removed (requested for real-world application), PaSTA-NS collapses to PaSTA at such strong autocorrelation, controlling FPR well yet traded-off with a decreased capability of accounting for data nonstationarity.

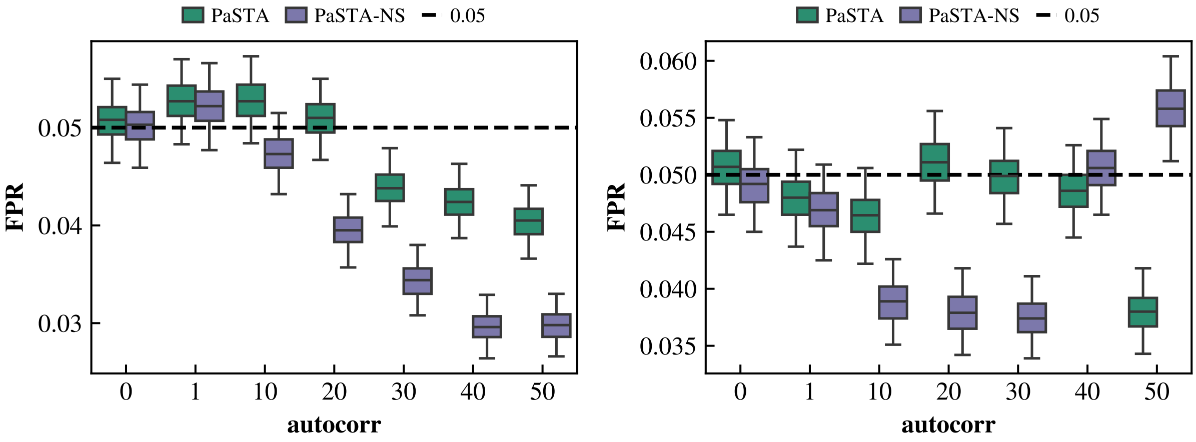


*Figure S3. FPR of PaSTA and PaSTA-NS.* Same as Fig. 4 but evaluated with 10,000 simulated map pairs per autocorrelation strengths.

#### Evaluating empirical brain maps with spatial trends

Fig. 5 compares PaSTA and benchmarks in assessing pairwise associations between empirical brain maps with spatial trends removed. Here, we also compare the statistical significance derived by PaSTA and benchmarks between raw brain maps with large-scale spatial trends (Fig. S4). PaSTA-NS, PaSTA, spin test, eigenstrapping, and BrainSMASH reported 5, 5, 7, 6, 6 significant findings, respectively. Like Fig. 5 in the main, PaSTA is more conservative than established methods when assessing empirical brain maps. Note that results of PaSTA and PaSTA-NS are overall similar except comparisons involving the neurosynth map. This is because when large-scale spatial trends are interpreted as autocorrelation, the range of autocorrelation is too long to allow parcellations of data. PaSTA-NS only parcellates the neurosynth map into two regions for nonstationarity detection. However, it should be noted that spurious autocorrelation arising from trending confounds is not appropriately controlled when trends are modelled as autocorrelation. As such, mean (spatial trend) analysis is a prerequisite for reliable significance inference.

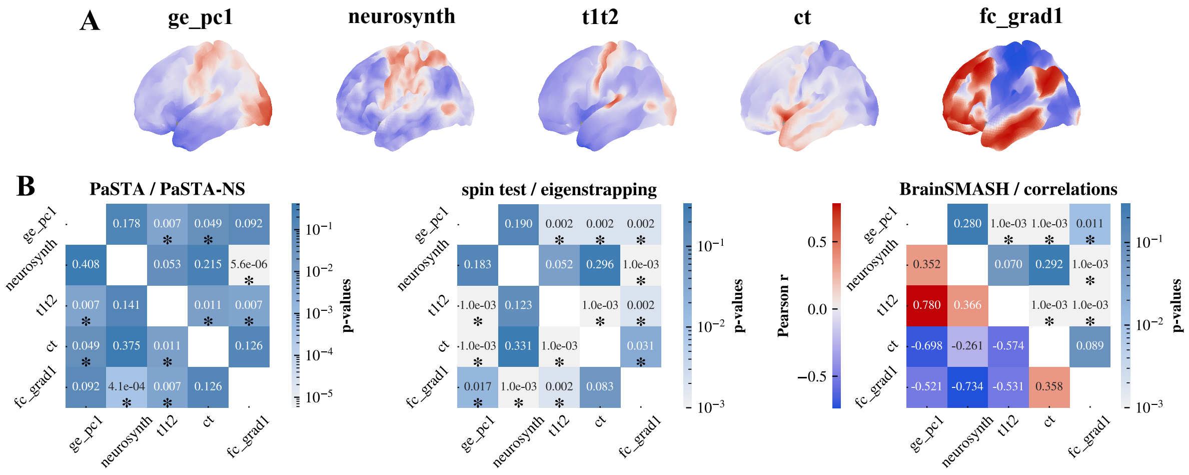


*Figure S4. Statistical significance of pairwise associations between empirical brain maps with spatial trends retained.* A) Visualization of brain maps with spatial trends. B) Pairwise associations between empirical brain maps. It displays significance p-values computed using PaSTA (left, upper triangle), PaSTA-NS (left, lower triangle), spin test (center, upper triangle), eigenstrapping (center, lower triangle), and BrainSMASH (right, upper triangle), and the Pearson correlation coefficient (right, lower triangle) for each pair of detrend maps. Significant p-values (<0.05) are marked with the asterisk symbol *.

### Effect of parameter selection in variogram estimation

Parameters $M$ and $qd$ (similar to $nh$ and $pv$ in BrainSMASH) are critical for accurate and reliable empirical variogram estimation, and therefore directly impact the efficacy in type-I error control. We suggest larger $M$ and $qd$ are typically beneficial and employ this strategy in PaSTA. Here, we detail the rationale for this choice.

We begin by introducing the basics of variogram. The top left panel in Fig. S5 demonstrates an example variogram. The semivariance increases as a function of lag distances, reaching the sill (i.e., sample variance, 1 in the example) at long distances where observations are spatially independent. The distance at which near-independence is observed is called range.

For empirical variogram estimation, the $qd$ parameter ($pv$ in BrainSMASH) determines the maximum lag distance evaluated (i.e., the range of x-axis of the variogram), where the $M$ parameter ($nh$ in BrainSMASH) determines the number of sampled points on the variogram (i.e., points observed on the curve). When $qd$ is small and empirical variograms are estimated for a lag distance shorter than the range of autocorrelation, the empirical estimation for autocorrelation between long-distance pairs is missing. When $M$ is small and empirical variograms are under-sampled before reaching to the sill, the description of autocorrelation at short-range distances can be inaccurate. Both can result in inaccurate and unreliable variogram estimation.

This is supported by our spherical mesh simulation where different combinations of $qd$ (between 0.1 and 1) and $M$ (25, 100, or 400) are selected in PaSTA (Fig. S5). When autocorrelation is strong (e.g., $l \geq 20$), small $qd$ leads to increased FPR in both large and small $M$, yet this is not evident in weak autocorrelation (e.g., $l \leq 10$). This is because small $qd$ does not provide enough variogram coverage in lag distances. However, large $qd$ can also increase FPR when a small $M$ is selected (e.g., $l \geq 20$, $M = 25$). This is because a small $M$ restricts the number of lag distances evaluated before variogram reaching to the sill, resulting in inaccurate estimation particularly for short-range, strongly autocorrelated data pairs. In contrast, the combination of large $qd$ and $M$ provides a broad coverage of lag distances evaluated, robustly estimates the variogram and false positives. This observation in theory also applies to BrainSMASH, as well as to eigenstrapping when variogram is visualized to assess surrogates (Burt et al., 2020; Koussis et al., 2025).

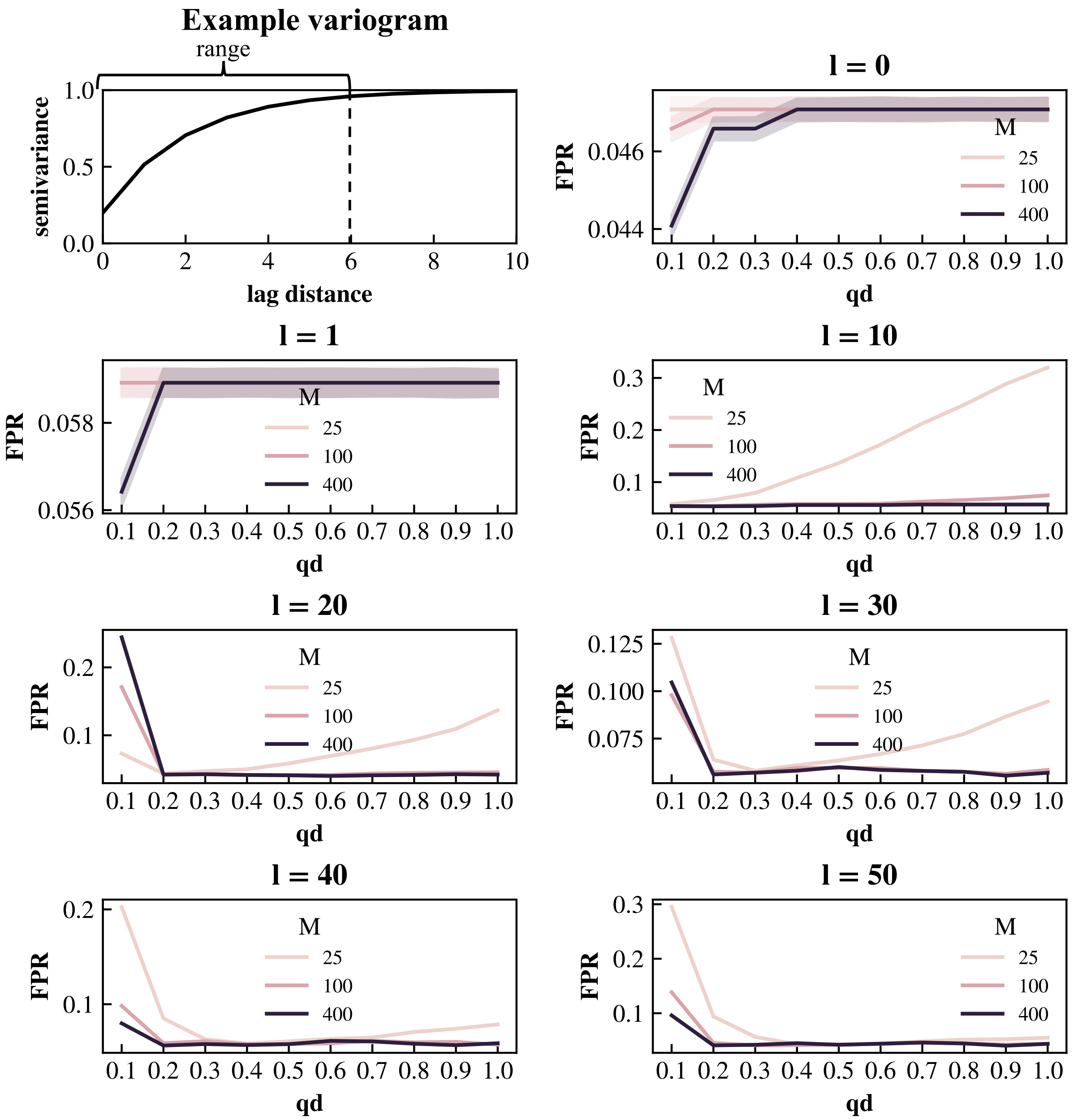


*Figure S5. Effect of parameter choices on FPR control.* Top left visualizes an example variogram. Other panels display the FPR achieved by PaSTA at various combinations of autocorrelation $l$ and parameters $qd$ and $M$. FPR are assessed with 1,000 independent map pairs per $l$.

**References**

Burt, J. B., Helmer, M., Shinn, M., Anticevic, A., & Murray, J. D. (2020). Generative modeling of brain maps with spatial autocorrelation. *NeuroImage*, *220*, 117038. https://doi.org/10.1016/j.neuroimage.2020.117038

Koussis, N. C., Pang, J. C., Phogat, R., Jeganathan, J., Paton, B., Fornito, A., Robinson, P. A., Misic, B., & Breakspear, M. (2025). Generation of surrogate brain maps preserving spatial autocorrelation through random rotation of geometric eigenmodes. *Imaging Neuroscience*, *3*, IMAG.a.71. https://doi.org/10.1162/IMAG.a.71
